# Can ancient DNA and other forms of time-sampled data aid in the inference of negative frequency-dependent selection?

**DOI:** 10.1101/2025.05.24.655935

**Authors:** Vivak Soni

## Abstract

Negative frequency-dependent selection (NFDS) is commonly viewed as the most efficacious form of balancing selection. Despite this, inferring NFDS remains challenging, and questions remain as to its relative importance in maintaining genetic variation in populations. Recent advances in both sequencing and genotyping technologies have resulted in a considerable increase in the number of publicly available human ancient DNA datasets, creating new opportunities for development of methods for the inference of NFDS from time-sampled data. In this perspective, I present three brief simulation studies to show how time-sampled data can aid improve inference power. First, I show how multiple time points can help us distinguish between recent NFDS and partial selective sweeps, as well as other forms of balancing selection, based on allele frequency trajectories. I then demonstrate how selective effects can be distinguished from population history based on changes in genetic variation and the site frequency spectrum over time.

Finally, I apply an approximate Bayesian computation approach to compare the power of multiple and single time point datasets in estimating the time for which NFDS has been shaping variation. Thus, I argue that data from multiple timepoints can facilitate the generation of new methodological approaches for better inference of NFDS.

## Introduction

Before advances in genomics allowed direct measurement of genetic variation across populations, a key debate in population genetics revolved around whether such variation was expected to be minimal or substantial. This controversy, known as the *classical/balanced debate* [1,2], largely focused on the role of selection—either purifying selection reducing variation or balancing selection preserving it [3].

However, even as next-generation sequencing later revealed abundant polymorphism in natural populations, the explanation shifted away from widespread balancing selection. Instead, the Neutral Theory of Molecular Evolution [4,5] gained prominence, proposing that much of the observed variation consisted of neutral alleles drifting randomly toward fixation or loss. This view has since been widely supported by empirical evidence [6]. Today, the extent to which balancing selection maintains genetic variation remains an open question [7,8], due in part to the challenges in detection, particularly on timescales that are neither extremely recent nor extremely long term (i.e. >25*N* generations, where *N* is the population size [9]). Indeed, estimates of the number of sites experiencing balancing selection have been limited to functional regions and are subject to a high level of uncertainty [10,11].

Despite evidence suggesting its role is more limited than once thought, balancing selection has been linked to a number of sites implicated in important functions, including sex determination [12], plant self-incompatibility [13–15], and the major histocompatibility complex (MHC) in vertebrates [16–18].

The term balancing selection encapsulates a number of selective mechanisms that maintain genetic variation, including negative frequency-dependent selection (NFDS), overdominance (also known as heterozygote advantage, whereby the heterozygous genotype has a higher fitness than either homozygous genotype) and spatial or temporal heterogeneity (whereby the fitness effect of a variant varies with environment and/or time). Of these, NFDS - the process by which the relative fitness of a variant is inversely proportional to its frequency in a population ([19], and see [20,21] for the first analytical treatments) – is commonly viewed as the most efficacious selective process maintaining balanced polymorphisms [22–27]. NFDS preserves genetic polymorphism in populations by favouring rare variants, which gain a selective advantage over common ones. Thus, rare alleles increase in frequency and persist rather than being lost from the population.

Although NFDS has been observed experimentally (e.g.[28.29]) and from analyses of phenotypic changes (e.g. [30–32]), distinguishing NFDS from other forms of balancing selection using population genomic data remains a challenge. This is because multiple forms of balancing selection leave similar genomic signatures [18] over differing timescales. The initial phase following the introduction of the selected mutation involves a rapid increase in frequency over time in the population, as the selected allele sweeps to its equilibrium frequency. Such partial sweep patterns can result in an excess of intermediate frequency alleles and extended linkage disequilibrium (LD – the correlation among alleles from different loci), as well as extended haplotype structure ([33–36], and see reviews of [37,38]) and weaker genetic structure [39]. These signals are fleeting however, as recombination breaks up haplotype and LD patterns, potentially resulting in reduced detection power [9,12,40,41] (and see reviews of 42,43]). At this point there is what Soni and Jensen [9] describe as a “large temporal gap”, before there is power to detect balancing selection using site frequency spectrum (SFS) based approaches. It is only once the balanced mutation has been segregating for considerable evolutionary time (>25*N* generations) that we have strong power to detect NFDS using SFS-based methods, as new mutations have accrued on the balanced haplotype, resulting in a skew in the SFS toward intermediate frequency alleles and an excess of variation in the neighbourhood of the selected locus. Finally, trans-species polymorphisms can be an informative signal of NFDS in species whose expected coalescent time is predated by their divergence time [44,45].

Despite these numerous signatures, a number of other neutral and selective processes can result in highly similar genomic patterns and thereby confound inference (see Table 1 of [42]). Thus, given the temporal gap, the confounding effects of other population genetic processes, and that observed levels of variation in numerous populations have been explained without positive or balancing selection (e.g. [46–50]), inference of NFDS remains challenging. However, time-sampled datasets in which we have access to genomic information over multiple time points - not just the present day – have the potential to improve inference of NFDS and other forms of balancing selection. Although time-sampled data of viral genomes has been available for some time (e.g.[51–53]), recent advances in both sequencing and genotyping technologies, as well as in protocols for handling degraded DNA from archaeological material (also known as ancient DNA, or aDNA), has resulted in a considerable increase in the number of publicly available human aDNA datasets, as exemplified by the Allen Ancient DNA Resource (AADR), a database of 12,761 ancient genomes, ranging across 10 time points, from hunter gatherers through to antiquity [54]. This glut of new data creates new opportunities for the development of methods for the inference of NFDS. In this perspective I explore these opportunities, identifying signals of NFDS that can be inferred across multiple time points. I demonstrate the potential increase in inference power across multiple time points via simulation, which will hopefully yield fruitful avenues for future studies on empirical time-sampled data.

## Inferring recent NFDS from population genomic data

Three challenges exist when attempting to infer very recent NFDS. The first is that NFDS is likely often a transient process because this form of selection favours rare alleles, and therefore other evolutionary processes can result in strong fluctuations in allele frequencies and the loss of allelic lineages. For example, Ejsmond and Radwan [55] showed via simulation that population bottlenecks can result in a severe and rapid loss of variation in genes evolving under NFDS. Thus, this form of balancing selection can be fleeting due to the increased probability of stochastic loss of the selected loci at low allele frequencies. The second challenge when detecting NFDS is that the characteristic SFS-based signatures of NFDS – increased variation and a skew in the SFS toward intermediate frequency alleles – are absent until the balanced mutation has been segregating for long enough for the balanced haplotype to accrue variation (>25*N* generations [9]). A mutation under NFDS that has escaped stochastic loss follows the initial trajectory of a partial selective sweep, and distinguishing between these two selective processes – and indeed other forms of balancing selection - is the third challenge when inferring very recent NFDS. A newly introduced beneficial mutation will rapidly increase in frequency, and linked variation may increase in frequency along with it [56]. This process will increase LD and therefore a number of tests of selection have been developed in order to detect this pattern of increased non-random association between alleles (e.g. the *EHH* [ extended haplotype homozygosity –[33]] and *iHS* [integrated haplotype score – [35]] statistics). Though this beneficial trajectory will be affected by details such as the strength of selection acting on the beneficial mutation, the general trajectories are equivalent for selective sweeps, NFDS and overdominance, at least until the balanced mutation reaches its equilibrium frequency, meaning that NFDS and partial sweeps of positive selection cannot be easily distinguished from one another [12,57].

One avenue for addressing this problem is through tracking temporal changes in allele frequencies, and thus distinguishing very recent NFDS from other forms of balancing selection, as well as partial sweeps of positive selection. More specifically, allele frequency information from multiple time points might enable researchers to examine how the beneficial mutation trajectories between these different forms of selection diverge once the balanced mutation reaches its equilibrium frequency. To demonstrate this approach, I ran simulations in SLiM v4.2.3 [58] of a single equilibrium population, with a beneficial mutation occurring on a neutral background, across 100 simulation replicates. I simulated two selective sweep regimes, with population scaled strengths of selection of 2*Ns* = 100 and 1,000 (where *N* is the population size of 500 individuals, and *s* is the selective advantage of the mutant allele relative to the wildtype); a single overdominance regime in which the balanced mutation had a population scaled strength of selection of 2*Ns* = 100 and a dominance coefficient, *h* = 20 (see Supplementary Figure S1 for comparison with *h* = 1.5); and a single NFDS regime in which the mutation under NFDS had an equilibrium frequency, *F_eq_* = 0.5 (see Supplementary Figure S2 for comparisons with *F_eq_* = 0.1 and 0.25). I also modelled spatial selection in which two equilibrium populations were simulated, with a beneficial mutation introduced in one population. This mutation was deleterious the other population, with gene flow occurring between the two populations at a rate 4*Nm* = 5, where *m* is the migration rate (see the Methods section for details of simulation framework). For all models, I sampled the population every five generations, starting from the introduction of the beneficial allele, until it had been segregating for 1*N* generations. Figure 1a shows the mean frequency of the beneficial mutations across the 100 simulation replicates, whilst Figure 1b and 1c show mean Tajima’s *D* [59] and mean haplotype diversity respectively. These summary statistics were calculated across sliding windows of size 10kb, with a step size of 5kb. From Figure 1a it is clearly visible where the beneficial mutation trajectories for selective sweeps diverge from those of the mutations under various forms of balancing selection. Whilst the allele under NFDS sweeps to its equilibrium frequency and then fluctuates about that frequency (0.5 here), the selective sweeps continue to increase in frequency toward fixation. This process occurs more rapidly at the higher strength of selection (2*Ns* = 1,000), though it is notable that the allele under NFDS also initially increases in frequency rapidly, due to the extremely strong selection acting on it at very low frequencies. As its frequency increases, the rate of frequency change slows down. Similarly, overdominance follows a partial sweep trajectory to some intermediate frequency (determined by the strength of selection acting on the mutation, as well as the dominance coefficient). At this point however, the mutation maintains a much more stable frequency relative to NFDS, as the heterozygous genotype maintains a higher fitness relative to either homozygous genotype (whereas the frequency and fitness effect of the mutation under NFDS continue to oscillate). Spatial selection also follows a similar pattern to NFDS, whereby the initial sweeping phase is followed by a fluctuation in frequency of the balanced mutation that is determined by the migration rate. Indeed, for all models discussed, the rate of allele frequency change will be determined by the parameterizations, as evidenced by the differing trajectories in Supplementary Figures S1-S3. Importantly however, the shape of the trajectory remains relatively consistent across parameterizations, which is necessary for distinguishing between these different models of selection.

**Figure 1:**
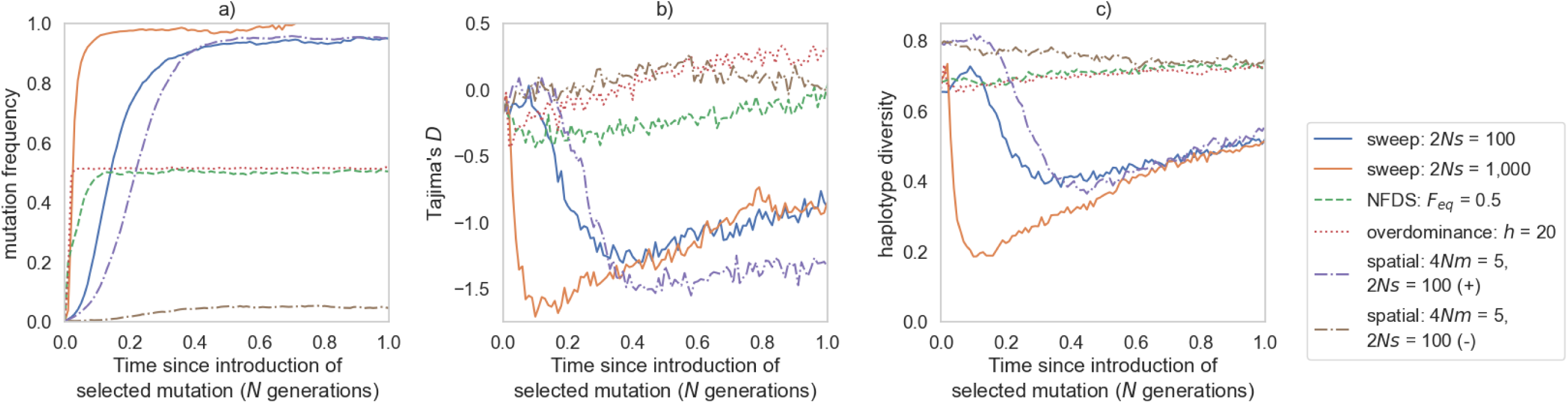
a) Mutation trajectories, **b)** Tajima’s *D* and **c)** haplotype diversity for selective sweeps, NFDS, overdominance in an equilibrium population, on a neutral background. For spatial selection, two equilibrium populations were simulated, with gene flow occurring between them. Summary statistics were calculated across sliding 10kb windows, with a 5kb step size. Only the focal window containing the mutation under selection is shown. All statistics shown are averages across 100 simulation replicates. Equilibrium frequency, *F_eq_* of the mutation under NFDS (green dashed line) is 0.5. The population scaled strength of selection, *2Ns* of selective sweeps, the overdominant mutation, and the mutation under spatial selection are 100. Selective sweeps were also simulated under a 2*Ns* value of 1,000 (orange line), where *N* is the population size of 500 individuals, and *s* is the selective advantage of the mutant allele relative to the wildtype. For spatial simulations, the migration rates is 4*Nm* = 5, where *m* is the migration rate. The purple dash-dotted line represents the population in which the balanced mutation is beneficial (marked by a + on the figure legend), whilst the olive dash-dotted line represents the population in which the balanced mutation is deleterious (marked by a - on the figure legend).

We can see the genomic signatures of these different selective regimes in Figures 1b and 1c. There is an initial increase in Tajima’s *D* around the selected locus immediately after the introduction of the beneficial mutation. A more positive Tajima’s *D* is caused by a skew in the SFS toward intermediate frequency alleles. Once a beneficial mutation has escaped stochastic loss, it will immediately increase in frequency, taking linked neutral variation with it, causing this resultant skew in the SFS. Likewise, there is an initial increase in haplotype diversity with this increase in allele frequencies, as shown in Figure 1c (except in the case of spatial selection, which is discussed further below). These patterns diverge as the allele under NFDS or overdominance reaches its equilibrium frequency. The continuing increase in frequency of the selective sweeps and their linked variation results in an increase in rare and high frequency alleles, and thus skews the SFS toward rare alleles, as evidenced by the reduction in Tajima’s *D*. Haplotype diversity is much reduced too, as the selective sweeps remove variation. By contrast, the balanced haplotype under NFDS maintains variation at the equilibrium frequency, and thus both Tajima’s *D* and haplotype diversity remain relatively consistent whilst the balanced mutation is segregating (though Tajima’s *D* notably fluctuates as the balanced mutation’s frequency oscillates around the equilibrium frequency). Indeed, both Tajima’s *D* and haplotype diversity will increase as the balanced mutation segregates for longer, as new mutations accrue on the balanced haplotype (see Figure 1 of [9]). A similar pattern of haplotype diversity occurs under overdominance, though the rate of increase in Tajima’s *D* is notably higher than under NFDS, which is explained by the consistent fitness advantage of the heterozygous genotype, allowing neutral variation to accrue on the balanced haplotype and thus skew the SFS toward intermediate frequency alleles. The increase in Tajima’s *D* under NFDS is tempered by the changing fitness of the balanced mutation, dependent on its frequency within the population. This increased rate of change in Tajima’s *D* under overdominance relative to NFDS is an important signature that may help distinguish between these two forms of balancing selection, and one that necessarily requires data from multiple time points.

Finally, the genomic signatures of spatial selection are distinct in the two populations, depending on whether the balanced mutation is beneficial or deleterious. If the balanced mutation is under strong selection, the trajectories of Tajima’s *D* and haplotype diversity be similar to those of selective sweeps where the mutation is positively selected for, albeit without reaching fixation. Where the mutation is deleterious, haplotype diversity is decreased (as expected under purifying selection). In figure 2b we see that Tajima’s *D* actually increases in this scenario as time since the introduction of the balanced mutation increases. Although purifying and background selection (BGS) result in a skew in the SFS toward rare alleles (and thus a reduction in Tajima’s *D*), migration will have the opposite effect, emphasizing the importance of modelling population history and gene flow when attempting to infer selective processes. Importantly, these signatures of spatial selection can be distinguished from NFDS, because in the latter case both Tajima’s *D* and haplotype diversity gradually increase with segregation time of the balanced mutation. Thus, if we have access to population genomic data that spans the initial phase of the beneficial mutation once it has escaped stochastic loss, we may be able to distinguish between NFDS and other forms of selection.

**Figure 2:**
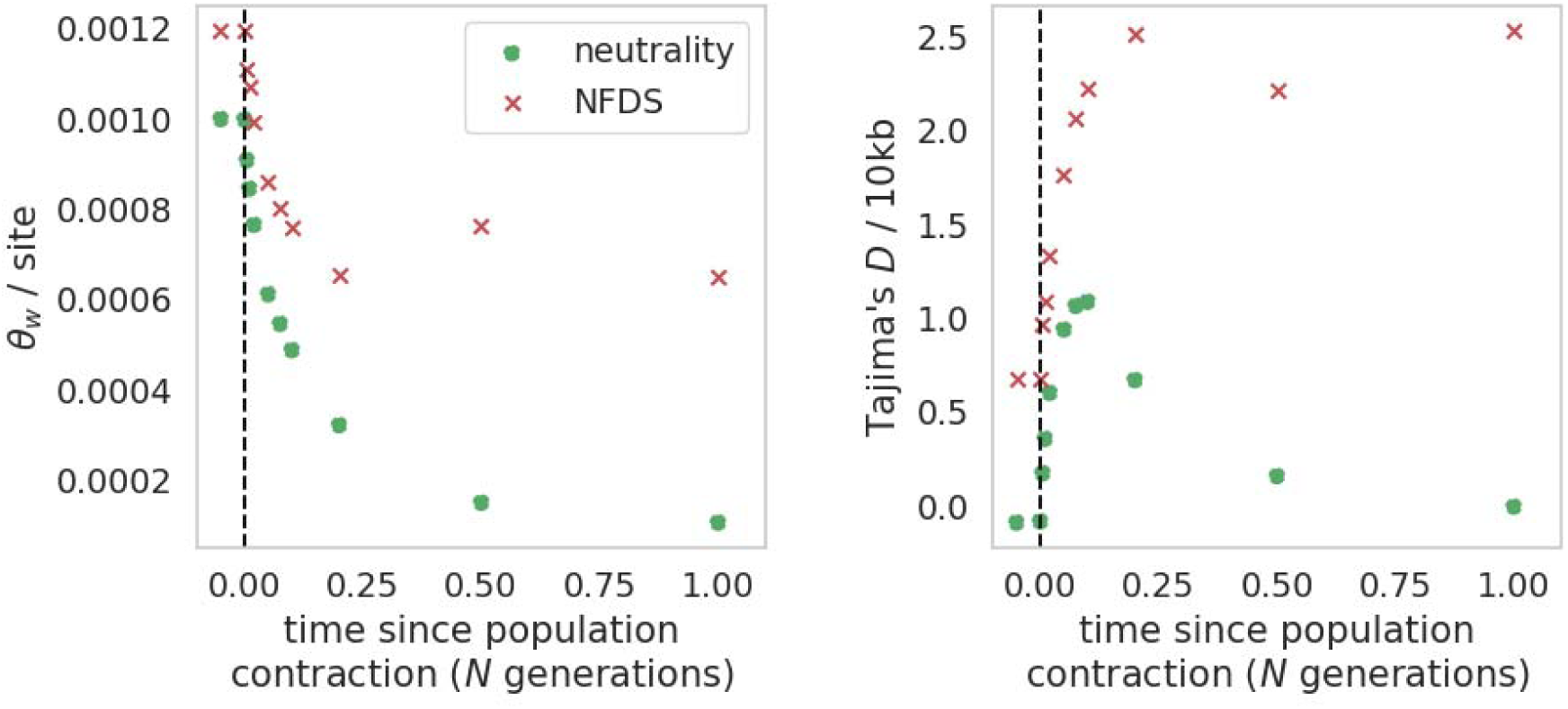
and Tajima’s *D* for neutral (green “o”s) and NFDS (red “x”’s) simulations in which an instantaneous population contraction occurred (black dashed line represents time of contraction). Population is reduced to 0.1*N_ancestral_* individuals, where *N_ancestral_* = 10,000. Data points to left of 0 on x axis indicate sampling prior to the population contraction. Equilibrium frequency, *F_eq_* of the mutation under NFDS is 0.5. Mutation under NFDS has been segregating for 75*N_ancestral_* generations at time of population contraction. Summary statistics were calculated across 10kb windows, then averaged across all windows and all 100 simulation replicates.

It is important to note that these processes may well leave similar signatures under certain parameterizations. Supplementary Figures S1-S3 depict mutation frequencies and genomic signatures under differing parameterisations (specifically, NFDS where *F_eq_* = 0.1 and 0.25 [S1]; overdominance where *h* = 1.5 [S2] and spatial selection where 2*N_e_s* = 10 [S3]). In the particular case of NFDS, we see that Tajima’s *D* does not appear to increase with segregation time under equilibrium frequencies lower than 0.5. This is due to the greater fluctuation in allele frequency occurring and thus less mutations accruing at intermediate frequencies on the balanced haplotype. Furthermore, one cannot simply apply these summary statistics to time-sampled data and identify recent NFDS. For instance, although the temporal evolution of the SFS helps us differentiate between selective sweeps and balancing selection, we know that there is little power to identify very recent balancing selection from the SFS alone [9] (and see reviews of [42,43]). Methods for detecting recent NFDS from time-sampled data would therefore need to harness linkage and haplotype information to identify candidate regions, and only then can allele frequency information be utilized for distinguishing between partial sweeps and NFDS. It is also important to model the underlying population history, as well as patterns of recombination and mutation, to avoid the confounding effects of these processes on inference of selection. With careful use of multiple aspects of population genomic data, there is considerable potential for inferring recent NFDS from time-sampled datasets, and distinguishing this form of balancing selection from other selective processes.

## Utilizing time-sampled data to account for the biasing effects of population history in NFDS inference

It is well documented that population history may confound the inference of selective processes (e.g. [60–71]). Soni and Jensen [9] showed how a reduction in population size can confound inference of balancing selection due to the skew in the SFS toward intermediate frequency alleles that both processes induce. Concurrently, many segregating variants might become fixed or lost, potentially breaking up the balanced haplotype. To account for population history – as well as other constantly acting evolutionary processes such as purifying and BGS effects, and mutation and recombination rate heterogeneity, Johri et al. [72,73] proposed the necessity of an evolutionary baseline model incorporating these processes prior to inferring comparatively rare or episodic processes such as positive and balancing selection. However, even with such baseline models, Soni and Jensen [9], have shown that power to infer balancing selection will be reduced under certain population histories.

To improve inference power when inferring NFDS under non-equilibrium population histories, we might attempt to perform genome scans across multiple time points, looking for consistent patterns across time. SFS-based inference approaches (e.g. [10,74]) detect the skew in the SFS generated by balancing selection, and the changes in the amount of genetic variation. If this signal holds in a genomic window across multiple time points, this might provide additional support for balancing selection. To demonstrate how these genomic signals might be maintained across multiple time points, I simulated a neutral region in a single population under an instantaneous population contraction, such that the population size was reduced to 10% of its initial size. I sampled this population multiple times, both prior to the contraction, and up to 1*N_ancestral_* generations after the population size change, where *N_ancestral_* was the initial population size of 10,000 individuals. I also simulated a functional region, modelling purifying and BGS, in which a single mutation under NFDS was introduced 75*N_ancestral_* generations prior to the population contraction (i.e. such that the balanced mutation had been segregating for long enough for SFS-based methods to have considerable power to detect NFDS). This population was sampled at the same timepoints as the neutral region simulations. I then calculated 0_W_ and Tajima’s *D* (Tajima 1989) across 10kb sliding windows with a 5kb step size for each time point. For full details of the simulation set up, see the Methods and Materials section. Figure 2 shows the changes in 0_W_ and Tajima’s *D* across time, both under neutrality, and for the functional region. NFDS elevates both statistics, as it increases the level of variation and the relative number of intermediate frequency alleles due to rare and high frequency alleles being lost or fixed respectively. The population contraction results in a reduction in 0_W_ as variation is lost from the population. The smaller population does not recover to the levels of variation prior to the size change. However, NFDS maintains a level of variation elevated above that of the neutral expectation. Conversely, the elevation in Tajima’s *D* due to the population contraction is lost after enough time has passed under neutrality, as the number of rare and high frequency alleles increase in the population. However, this is not the case under NFDS, where the elevation in Tajima’s *D* is maintained due to mutations on the balanced haplotype being maintained at intermediate frequencies.

Thus, we would expect to see consistent patterns across time points in genome scans, providing increased support for candidate regions under NFDS.

It is important to emphasize the need for an evolutionary baseline model [72,73] to generate a “null” expectation. The example given here would be relevant for humans and other species with coding-sparse genomes, where we have access to non-functional regions distant enough from any coding region to avoid the biasing effects of purifying and BGS [75], thereby modelling neutral processes independently of selective effects. With a population history inferred on these non-functional regions, we can infer the DFE of new mutations on functional regions, conditional on the demography already inferred. Other species such as viruses – in which the entire genome may be functional – would necessitate jointly modelling neutral and selective effects [46]. Once we have our baseline model, we can simulate under this model (sampling at the same time points that we have in our empirical data) and perform inference of NFDS on this model to infer null thresholds for inference, reasoning that these are the most extreme values of our statistic of interest that we can generate in the absence of NFDS. This process would be necessary for each sampled time point, mirroring our empirical data. Thus, if we identify regions that exceed these values in our empirical data, these are potential candidates for NFDS, and this can be verified across multiple time points.

## Inferring the segregation time of a mutation under NFDS

A parameter of interest when performing inference for NFDS is how long the balanced mutation has been segregating for (hereafter *T_b_*), because this parameter provides information on the duration of time for which NFDS has been shaping variation. However, we are generally limited to broad temporal categories based on the detectable signature. For example, Fijarczyk and Babik [42] categorize recent balancing selection as that in which the balanced mutation has been segregating for <0.4*N_e_* generations, whilst intermediate balancing selection is 0.4-4*N_e_* generations old, and ancient balancing selection is >4*N_e_* generations old. Bitarello et al. [43] provide a different but generally compatible schema, with recent balancing selection being <4*N_e_* generations old, long-term balancing selection >4*N_e_* generations old, and ultra long-term balancing selection being 4*N_e_* + *T_div_* generations ago (i.e. longer than the expected coalescence time between lineages present in species that diverged *T_div_* generations ago). Because different ages of selected alleles generate distinct signatures, we are able to infer the timescale on which balancing selection has been acting. A question then remains as to whether we can narrow down this temporal range further.

To answer this question, I used an approximate Bayesian computation (ABC) approach to attempt to infer *T_b_* from simulated data. Although there are numerous viable statistical frameworks for addressing such a question (including maximum likelihood and machine learning based approaches), ABC combines the benefits of Bayesian inference with the computational efficiency of working with summary statistics. Indeed, the simulation step removes nuisance parameters which are important confounders in population genetic inference. I ran simulations in which a mutation under NFDS had been segregating for 10*N_ancestral_* generations, with the aim of comparing inference power for a single time point, and multiple time points. I sampled the data at the end of this segregation time (which would be the only time point for the single time point analysis), as well as 0.02*N_ancestral_* generations prior, with the two time points together forming the multi time point data. This ancient time point was chosen as the 0.*02N_ancestral_* time point falls within the AADR aDNA dataset [54] and is therefore a reasonable timescale for which we have population genomic data in humans (and given that this database does not contain populations with multiple sampled individuals from the same time points and geographic locations, I limited the analysis to just two time points in total). This “empirical” data (i.e. the data that ABC would be used to fit a model to) was generated for four different population histories: equilibrium (no population size change); expansion (where the population size instantaneously doubled); contraction (where the population size instantaneously halved); and severe contraction (where the population size was instantaneously reduced by 90%). The Methods and Materials section contains details of how this data was generated. For ABC inference, 3,100 total values of *T_b_* were drawn from a uniform distribution with bounds of 0 and 100 *N_ancesltral_* generations and simulated in SLiM v4.2.3 [58] (see Methods section for details of the sampling scheme). Summary statistics were calculated across 10kb windows with a 5kb step size, with the mean and standard deviation for each window utilized as a summary statistic for ABC inference. Although any number of summary statistics might be used for inference with ABC, I used Tajima’s *D* [59], 0_W_, haplotype diversity, mean *r^2^*, and the number of singletons, given that these statistics together capture different aspects of the data. This created a total of 60 statistics for ABC inference from the single time point data, and 120 statistics for the multiple time point data.

Figure 3 shows the posterior distribution of *T_b_* for the four population histories, both for the single and multiple time point data. In each case, the inference based on a single time point results in a wide posterior distribution. By contrast, the posterior distributions from multiple time points are much more informative, with modal values either over or very close to the true value of *T_b_* of 10*N_ancestral_* generations. The increasing prevalence of time-sampled data may therefore facilitate more precise estimates of how long a mutation under NFDS has been segregating for.

**Figure 3:**
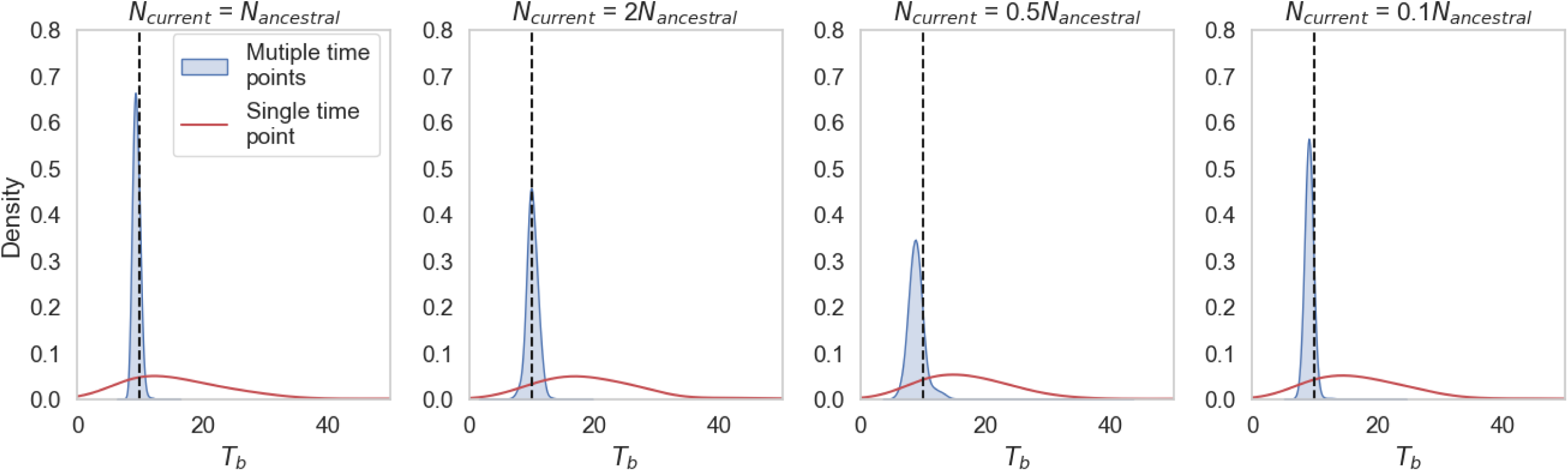
Posterior distributions from ABC inference of the segregation time, *T_b_* of the mutation under NFDS for four different population histories. Black dashed line represents the true value of *T_b_* (10*N_ancestral_* generations, where *N_ancestral_* is the initial population size of 10,000 individuals). Multiple data points (blue shaded distributions) include sampling at the current day, and 0.02*N_ancestral_* generations prior to current day,. Single time point data (red unshaded distributions) is for sampling at the current day only only. Four underlying population histories are represented. From left to right: Population equilibrium (*N_current_* = *N_ancestral_*), population expansion (*N_current_* = 2*N_ancestral_*), population contraction (*N_current_* = 0.5*N_ancestral_*), and severe population contraction (*N_current_* = 0.1*N_ancestral_*). All population size changes occurred instantaneously, 0.01*N_ancestral_* generations prior to the current day.

An important caveat here is that this analysis focuses on a functional region in which a mutation under NFDS is segregating, under the assumption that this functional region has already been identified as a candidate region for NFDS. It is also assumed that due diligence has been performed in terms of inferring an evolutionary baseline model (i.e. the demographic history, DFE, and mutation and recombination rates are known). With real world data, these processes will have to be inferred prior to inference of *T_b_*. It is encouraging however, that inference of *T_b_* from multiple time point data is accurate even when the mutation under NFDS has been segregating for just 10*N_ancestral_* generations, given that this timeframe sits within the temporal gap between 1*N* and 25*N* generations identified by Soni and Jensen [9]. The drawback of such ABC approaches is that they are computationally expensive, particularly when exploring a large parameter space and under complex population histories.

## Concluding thoughts

The simulation studies presented here illustrate the potential of time-sampled data to radically improve inference of NFDS, as well as other forms of balancing selection from population genomic data. Whereas the signature of a hard selective sweep is fleeting [76,77], balancing selection can persist across both short and long timescales. For examples, Laval et al. [78] found that the haemoglobin {J^S^ sickle mutation – a textbook case of natural selection maintaining a deleterious mutation at high frequencies via balancing selection – initially arose in humans ∼22,000 years ago. At the other extreme, Leffler et al. [45] found multiple instances of polymorphisms shared by humans and chimpanzees being maintained by balancing selection, indicating that the balanced mutation has been segregating at least since before the human chimpanzee split time 7-13 million years ago [79]. As I have shown in this study, time-sampled data may help researchers to more precisely date how long NFDS has been acting on a mutation, as well as increasing the amount of relevant information with which to tease apart neutral and selective processes.

As with any simulation study however, an important qualification is that this is “perfect” data with many assumptions that will not necessarily hold in natural populations. For example, I have assumed no mutation and recombination heterogeneity, which will impact inference of NFDS, and have modelled a single isolated population, when population structure is a well-known confounder of balancing selection inference. Indeed, hidden structure – where multiple populations are assumed to represent just a single population – results in a considerably higher false positive rate when performing balancing selection inference [9]. When handling aDNA there are a number of further complications to consider. Though DNA can preserve for hundreds of thousands of years under favorable conditions (e.g [80,81]), genetic material decays progressively through time following the death of an organism, due to cellular repair functions no long functioning postmortem [82,83].

Humidity, temperature, salinity, pH, and microbial growth all influence DNA preservation. The extent and nature of their effect on DNA preservation will vary depending on the archaeological site and stratum in which they are located (see reviews of [84,85]). A further problem is that of DNA contamination. Contamination can occur both from the presence of microbial and environmental sources [86], as well as human DNA introduced during sample extraction or laboratory processing [87–89]. Methods such as that of [89] have been developed to quantify the proportion of present-day DNA contamination in aDNA datasets, and thus any attempt to infer population genetic processes from aDNA datasets must model the effects of DNA damage and contamination where possible, and model the possible inference impact of the uncertainty due to aDNA damage by drawing error rates from an experimentally-informed range to account for the biasing effects such damage and contamination might have on downstream inference.

Another challenge that is specific to humans is the geographic mobility our species throughout much of our history. This mobility has meant that even if samples were of the same geographic location, the ancient sample may not represent the ancestral population of the current day sample. A potential solution is to attempt to connect the composition of ancestry in the ancient sample with the composition of ancestry in the modern sample, facilitating the comparison of allele frequencies in the mapped ancestry components of the genomes. Other forms of time-sampled data will come with their own challenges and potential. For example, frequency oscillations over time observed at the phenotypic level have previously been cited as evidence of NFDS [90–92]. These fluctuations at the ecological timescale are detectable in population genomic data in smaller populations, and time-sampled data might facilitate the tracking of these fluctuations in allele frequencies through time, bridging both phenotypic and genomic data, and ecological and evolutionary timescales. Such analyses may also prove viable with data from biodiversity collections (e.g. [93])

Thus, by constructing a baseline model that accounts for both the specific nature of the data, as well as the constantly occurring population genetic processes such as genetic drift, purifying and BGS, and mutation and recombination rate heterogeneity, we can harness the extra information from time-sampled data to distinguish NFDS from other selective and neutral processes, as well as quantify how long NFDS has been maintaining variation for. Because NFDS is constantly shaping variation (for example, increasing the skew in the SFS as new mutations accrue on the balanced haplotype), even a small number of closely grouped time points provide increased resolution into detection of this process. The increasing prevalence of aDNA and time-sampled datasets in other organisms can spur development of new methods for inference of NFDS and other population genetic processes.

## Methods and materials

### Generalized simulation framework

All simulations were run forward-in-time using SLiM v.4.2.3 [58], using parameters previously inferred in humans. 100 replicates were simulated for any given parameter combination or simulation regime. All simulations had an initial burn-in time of 10*N_ancestral_* generations, where *N_ancestral_*was the initial population size of 10,000 individuals [94] unless otherwise stated. A single population was simulated with a fixed mutation rate of 2.5 x 10^-8^ per base pair per generation [95], and a fixed recombination rate of 1 x 10^-8^ per base pair per generation [96]. Negative frequency-dependent selection was modelled such that the selection coefficient of the balanced mutation, *S_bp_*, was dependent on it’s frequency in the population: *S_bp_* = *F_eq_* - *F_bp_*, where *F_eq_* is the equilibrium frequency of the balanced mutation, and *F_bp_* is the frequency of the balanced mutation. The dominance coefficient, *h* of the balanced mutation under NFDS or spatial selection was 0.5. Simulation replicates in which the balanced mutation failed to establish by reaching a frequency of 0.1 were discarded and restarted.

### Selective sweep and NFDS trajectory simulations

A population of *N_ancestral_*= 500 individuals was simulated on a 100kb neutral background. This smaller population size was used for computational efficiency. A single beneficial mutation was introduced after the 10*N_ancestral_* burn-in. For selective sweep simulations, 2 different population scaled strengths of selection were modelled: 2*Ns* = 100 and 1,000, where *N* = *N_ancestral_* and *s* is the selective advantage of the mutant allele relative to the wildtype. Simulations in which the beneficial mutation failed to fix were discarded and restarted. For NFDS simulations, equilibrium frequencies of 0.5, 0.25, and 0.1 were modelled. For overdominance simulations, dominance coefficients of 20 and 0.1 were modelled. For spatial selection, two populations of size *N_ancestral_* = 500 individuals were modelled, with gene flow between them at a rate of 4*Nm* = 5, where *m* is the migration rate. The strength of selection acting on the introduced balanced mutation was 2*Ns* = 100, with a beneficial fitness effect in one population, and a deleterious effect in the other. For each simulation model, 20 individuals were sampled every 5 generations from the introduction of the beneficial mutation until either its fixation, or a further 10*N_ancestral_* generations had passed. In the case of spatial selection, each population was sampled.

### Population contraction simulations

A single population was simulated, undergoing an instantaneous population contraction 0.1*N_ancestral_* generations following the burn-in, reducing to 0.1*N_ancestral_* individuals. 20 individuals were sampled at 14 time points, with two prior to the population contraction, one at the time of contraction, and then at 11 time points post-size change, with the final time point 1*N_ancestral_* generations since the population contraction. Following the approach of Soni and Jensen [75], I separately simulated a 1Mb non-functional region, and a 32,657bp functional region. The functional region consisted of nine exons (of size 1,317bp) and eight introns (of size (1,520bp), flanked by intergenic regions of size 4,322bp. The numbers of introns and exons per functional region were obtained from Sakharkar et al. [97], with mean intron length taken from Hubé and Francastel [98]. Finally, the lengths of exons and intergenic regions were averages estimated from Ensembl’s GRCh38.p14 dataset [97], obtained from Ensembl release 107 [98]. Mutations in intronic and intergenic regions were all strictly neutral, whilst exonic mutations were drawn from the discrete distribution of fitness effects (DFE) inferred by Johri et al. [47] in humans. A single mutation under NFDS, with an equilibrium frequency, *F_eq_* of 0.5 was introduced into the middle exon following the burn-in period.

Tajima’s *D* [59] and and 0_W_ were calculated across 10kb sliding windows with a step size of 5kb, using the python implementation of libsequence (version 1.8.3, [101]), with mean values calculated across all windows and simulation replicates.

### Approximate Bayesian computation (ABC) inference

To generate simulated empirical data for ABC inference, a single population of size *N_ancestral_* = 10,000 individuals was simulated for 100 replicates. A single functional region was simulated, using the same genomic structure and DFE as outlined in the section titled “Population contraction simulations”. A mutation under NFDS was introduced immediately after the burn-in, and 10*N_ancestral_* generations prior to sampling. Four population histories were simulated, with ABC inference performed on each separately: population equilibrium (*N_current_* = *N_ancestral_*, where *N_current_* is the population size at time of sampling); population expansion (*N_current_* = 2*N_ancestral_*); population contraction (*N_current_* = 0.5*N_ancestral_*); and severe population contraction (*N_current_* = 0.1*N_ancestral_*). The instantaneous size change occurred 0.01*N_ancestral_* generations prior to sampling.

ABC inference was used to infer a single parameter of interest, *T_b_*, the segregation time of the mutation under NFDS. Values for this parameter were drawn from a uniform distribution with a range between 0*N_ancestral_* and 100*N_ancestral_*. A total of 3,000 different values of *T_b_* were simulated for 100 replicates, across three rounds of ABC. First, 1,000 values of *T_b_* were drawn from the uniform distribution, and simulated. ABC inference was performed on these simulations, generating a posterior distribution. A further 1,000 values were draw from this posterior distribution and ABC inference performed again. This process was repeated for a third time, giving a total of 3,100 simulated values of *T_b_*. The mean and standard deviation of the number of singletons, Tajima’s *D* [57], 0_W_, haplotype diversity, and mean *r^2^* were used for ABC inference, all calculated across 10kb sliding windows with a 5kb step size, using the python implementation of libsequence (version 1.8.3,[101]), with mean values calculated across all windows and simulation replicates, with each of the six genomic windows treated as a separate summary statistic. For the single time point inference, 20 individuals were sampled at the termination of the simulation, *T_b_* generations after the simulation burn-in, yielding a total of 60 summary statistics. For multiple time point inference, the population was sampled 0.02*N_ancestral_* generations prior to the termination of the simulation, as well as at its termination, yielding a total of 120 summary statistics.

The “neural net” regression method with the default parameters provided by the R package “abc” [102] was used to generate posterior distributions. A 100-fold cross validation analysis was performed in order to determine the performance and accuracy of inference for tolerance values of 0.05, 0.08, and 0.1, with a tolerance of 0.08 identified as the most accurate. This value was employed for inference of final parameter values, meaning that 8% of all simulations were accepted by the ABC to estimate the posterior probability of parameter estimates. Inference was performed 50 times, with the mean of the weighted medians of the posterior estimates taken to determine point estimate of *T_b_*.

## Data availability

All scripts to generate and analyze simulated data are available at the GitHub repository: https://github.com/vivaksoni/NFDS_time_sampled_data

## Acknowledgements

I am grateful to Jeffrey D. Jensen (Arizona State University) for valuable discussions relating to population genetic inference, and feedback on this manuscript.

