## Supplementary Figure for "Can ancient DNA and other forms of time-sampled data aid in the inference of negative frequency-dependent selection?"

**Supplementary Figure S1**

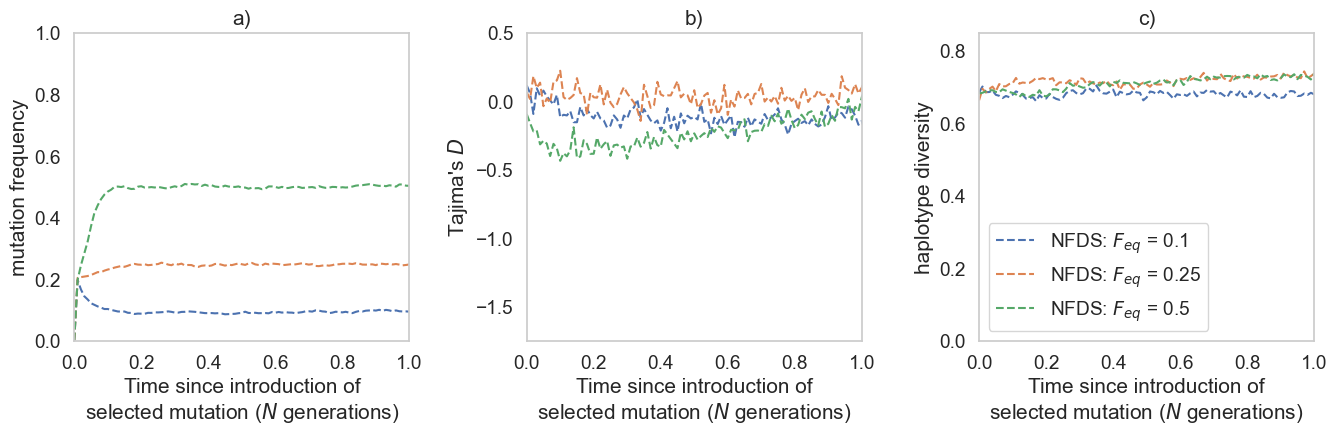

**S1: a)** Mutation trajectories, **b)** Tajima’s *D* and **c)** haplotype diversity for NFDS under different equilibrium frequencies (*F_eq_*). Summary statistics were calculated across sliding 10kb windows, with a 5kb step size. Only the focal window containing the mutation under selection is shown. All statistics shown are averages across 100 simulation replicates.

**Supplementary Figure S2**

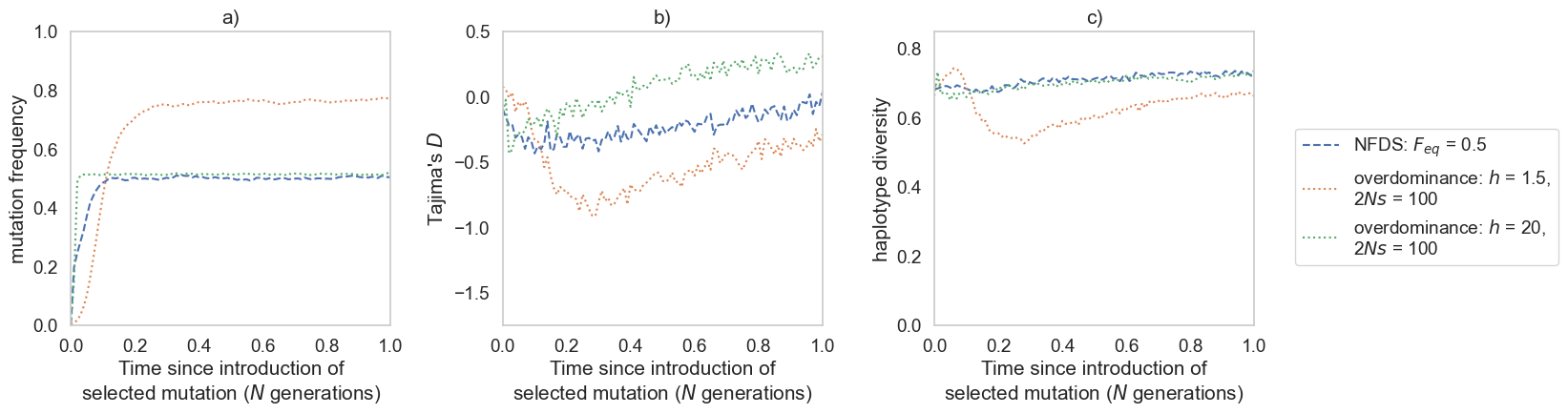

**S2: a)** Mutation trajectories, **b)** Tajima’s *D* and **c)** haplotype diversity for overdominance in an equilibrium population, on a neutral background. NFDS has been included for comparison. Summary statistics were calculated across sliding 10kb windows, with a 5kb step size. Only the focal window containing the mutation under selection is shown. All statistics shown are averages across 100 simulation replicates. Equilibrium frequency, *F_eq_* of the mutation under NFDS (green dashed line) is 0.5. The population scaled strength of selection, *2Ns* of the overdominant mutation is 100, where *N* is the population size of 500 individuals, and *s* is the selective advantage of the mutant allele relative to the wildtype.

**Supplementary Figure S3**

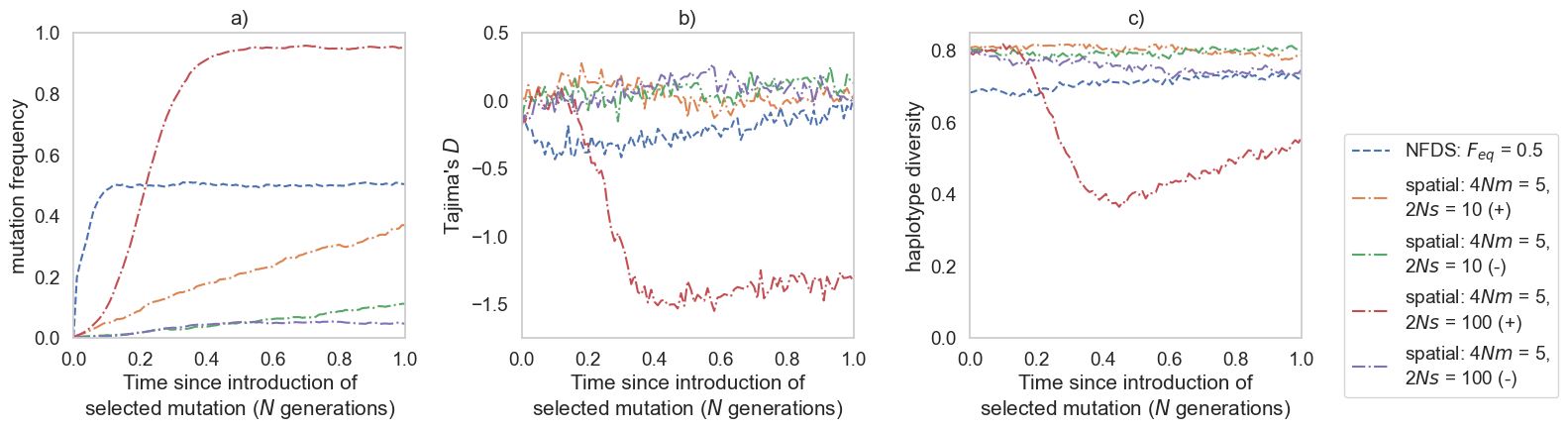

**S3: a)** Mutation trajectories, **b)** Tajima’s *D* and **c)** haplotype diversity for spatial selection, where two equilibrium populations were simulated, with gene flow occurring between them. NFDS has been included for comparison Summary statistics were calculated across sliding 10kb windows, with a 5kb step size. Only the focal window containing the mutation under selection is shown. All statistics shown are averages across 100 simulation replicates. Equilibrium frequency, *F_eq_* of the mutation under NFDS (green dashed line) is 0.5. The population scaled strength of selection, *2Ns* of the mutation under spatial selection is 10 or 100, where *N* is the population size of 500 individuals, and *s* is the selective advantage of the mutant allele relative to the wildtype. For spatial simulations, the migration rates is 4*Nm* = 5, where *m* is the migration rate. The red and orange dash-dotted lines represent the population in which the balanced mutation is beneficial (marked by a + on the figure legend), whilst the green and purple dash-dotted line represent the population in which the balanced mutation is deleterious (marked by a - on the figure legend).
